# How the brain distinguishes a bilingual’s languages is partly shared across people and partly their own

**DOI:** 10.64898/2026.09.18.752561

**Authors:** Sinem Demirkan, Demian Wassermann, Benoît Sagot

## Abstract

How personal is a cognitive distinction? People performing the same task show similar brain activity, which group-level neuroimaging builds on. Yet brain organization differs reliably between individuals, and people bring different experiences to every task. Bilingual language use is a natural test case. We asked whether the distinction between a bilingual’s languages is common to everyone performing the task, or whether each person keeps a part of their own. We decoded cortex-wide fMRI activity within each of 77 bilinguals naming pictures in Chinese, their dominant language, or English, switching languages from one moment to the next. Within a person, the language in use can be classified on single namings (balanced accuracy 59.6%, chance 50%). Classifiers trained on one person also transfer to others (56.9%). To locate what transfers, we estimated the main axes of trial-to-trial task activity in other participants, without language labels. Transfer runs entirely along these axes, and the first 50 alone match the full cortical pattern (57.2%). Without these axes, transfer between people nearly disappears, yet decoding within a person is unchanged. The remaining activity is stable enough across runs to identify most individuals (74%, chance 1.3%). Age of English acquisition is reflected along the shared axes, accounting for individual differences in transfer (cross-validated R2 = 0.19), but not in the remaining activity. The distinction between a bilingual’s languages is thus partly carried by axes common to the group, and partly by a stable signature specific to each person.

## Introduction

Brain function differs greatly from one individual to another. This has been demonstrated both in the intrinsic organization measured when people are at rest [1–3] and in the activity caused by cognitive tasks [4, 5]. These differences are stable for each individual and are thought to contribute to differences in behavior and cognition [1, 3, 6, 7]. Moreover, it is now possible to map out the functional organization of the brain both in large groups [8, 9] and within individuals [2, 7, 10]. These approaches, however, typically rely on large samples or on many hours of data from each person, often recorded at rest [1–3, 11]. Consequently, averaging signals across a large number of participants remains common in human neuroimaging today.

One field where individual-level approaches remain limited is bilingualism. Paradoxically, bilinguals are considered a very diverse group of individuals in the way they engage with language. Bilinguals differ in when and how they acquired their languages, how proficient they are in each, and how they use them [12, 13], and these differences are associated with variation in brain structure and function [13–15]. Yet neuroimaging studies of bilinguals typically collect a limited number of trials from each person and analyze the brain region by region at the group level. All bilingual speakers nevertheless move between languages and, at any given moment, use one of them. Selecting the appropriate language while managing the other is referred to as bilingual language control [16–18]. We refer to the state of the brain while using one language rather than the other as bilingual language state. It is not exclusively linguistic and includes any difference in how a picture is named in each language, from planning and control to articulation.

The two languages of a bilingual give rise to distinguishable patterns of neural activity. Earlier work identified differences in the strength of regional activity [19–22]. More recent work used multivariate decoding, which infers the language in use from patterns of brain activity [23], and found language information in fine-grained voxel patterns within regions or local neighborhoods [24, 25]. In both approaches, each region is examined separately. At the same time, language processing has been shown to engage a distributed fronto-temporo-parietal network that is similar across languages and speakers [26–28]. When one of these languages is acquired later in life, using it additionally recruits regions associated with language and cognitive control [18, 29, 30]. This means that the distinction between languages could exist in a more distributed pattern than what has been captured by the regional approaches.

In this work, we used cortex-wide multivariate modeling of brain activity in individual bilinguals. For this, we used an openly available dataset that is unique in bilingual neuroimaging in the combination it offers [31]. It combines a large sample of 77 Chinese–English bilinguals, for whom Chinese was the first and dominant language and English the second. Both languages of a bilingual are sampled trial by trial within the same task, and are intermixed so that each participant repeatedly switched between them. We fit linear models to the cortex-wide activity of each bilingual and tested them both within each person and across 5,852 pairs of participants, training a classifier on one person, the *teacher*, and testing it on another, the *learner* [32, 33]. We address three main questions: i) Is the distinction between languages expressed at the scale of the whole cortex, beyond the regional patterns described so far? ii) Is this distinction shared across individuals? and iii) Does the shared part lie along the main axes of task activity common to all participants, and is what remains noise or a signature specific to each person?

## Results

### Language state is expressed in a cortex-wide pattern

On each trial, participants saw a line drawing of an object and named it aloud in Chinese or English according to a color cue, with the two languages rapidly intermixed within each of two fMRI runs [31]. The dataset authors provided activity estimates for every naming trial by modeling one trial at a time as the event of interest [31]. We divided the cortex into the same 800 parcels in every participant [9] and averaged activity across voxels within each parcel (Fig. 1*A*). This gave each trial a cortex-wide pattern of 800 values, which we used to distinguish trials cued for Chinese from those cued for English. Any difference between naming a picture in one language and in the other, from planning and language control to articulation, could contribute to this distinction.

**Figure 1.**
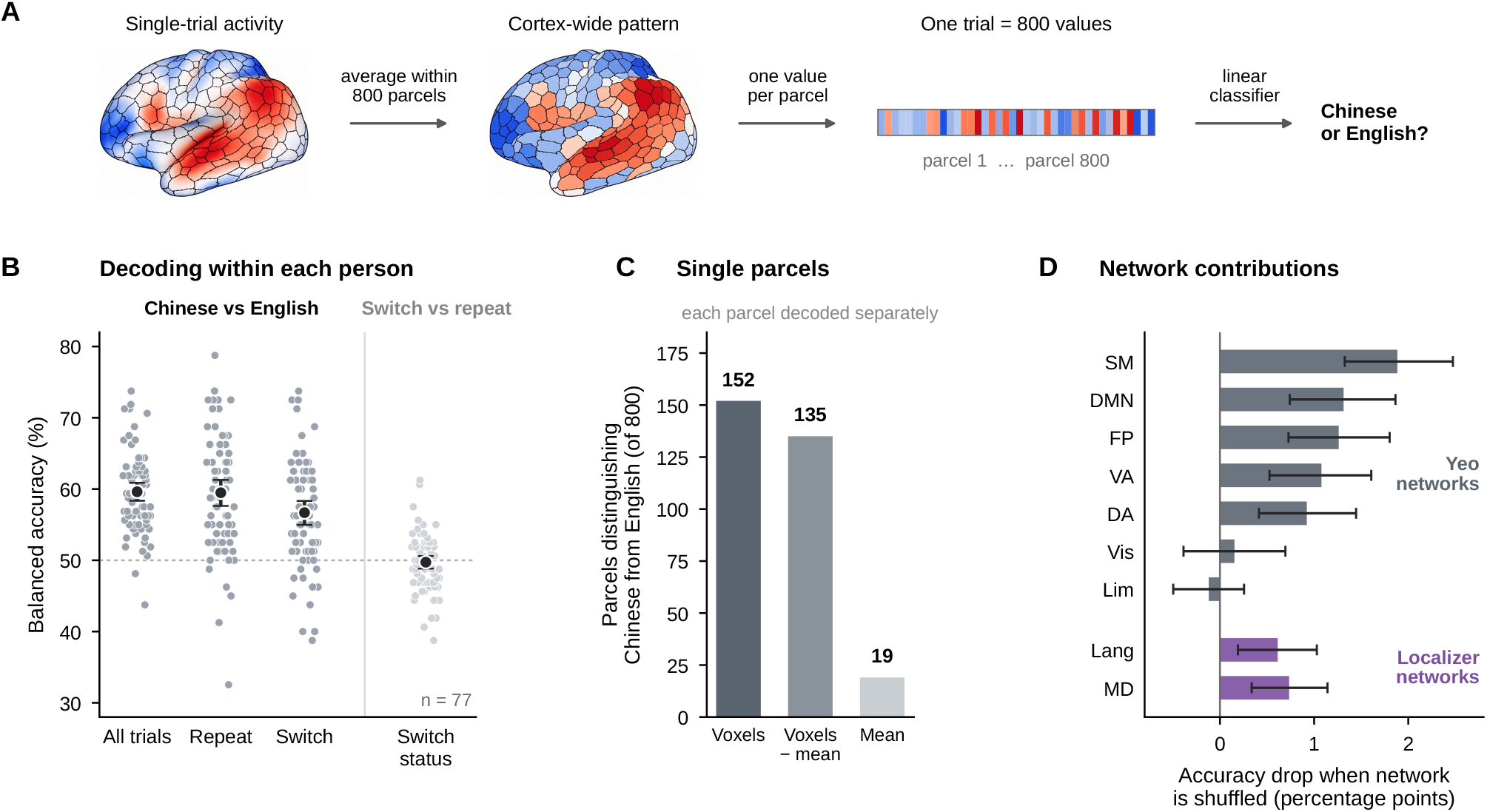
Language state is expressed in a cortex-wide pattern. (*A*) Single-trial activity is averaged within 800 parcels, and the resulting cortex-wide pattern of 800 values is used by a linear classifier to distinguish Chinese from English trials. (*B*) Decoding within each person across runs, for all trials, repeat trials and switch trials, and decoding of switch status from the same trials. Gray points show participants (*n* = 77); black points and error bars show the mean and 95% CI; the dotted line marks chance. (*C*) Parcels distinguishing the languages one at a time (group level, FDR-corrected) from voxel patterns, voxel patterns without the parcel mean, or the parcel mean alone. Parcel means rarely distinguish the languages one at a time, yet all 800 together do (*B*). (*D*) Permutation feature importance of the seven Yeo networks and of the language and multiple-demand parcel sets, as the drop in held-out balanced accuracy when a network’s PCA component scores are shuffled. SM, somatomotor; DMN, default; FP, frontoparietal (control); VA, ventral attention; DA, dorsal attention; Vis, visual; Lim, limbic; Lang, language; MD, multiple demand. All CIs are from a participant bootstrap (5,000 resamples).

To test if these patterns reliably distinguished language state, we trained a logistic regression classifier on patterns from one run and tested it on new trials from the same person’s other run. The classifier achieved a balanced accuracy of 59.6% across participants (95% confidence interval (CI) [58.4%, 60.9%]; range: 43.8%–73.8%; Fig. 1B). Whole-cortex decoding accuracy was not related to mean temporal signal-to-noise ratio (tSNR) across participants (*r* = *−*0.10, *p* = 0.40; Spearman *ρ* = *−*0.10, *p* = 0.38; *SI Appendix*, Fig. S1).

This distinction was robust to how cortical activity was represented. Changing the spatial granularity of the parcellation, or reducing it with PCA, had little effect on decoding performance (*SI Appendix*, Fig. S2).

Our cortex-wide representation was designed to capture language state at a different spatial scale from previous regional approaches. But was this truly a different scale of representation? We tested this by first examining if individual cortical parcels distinguished Chinese from English through fine-grained patterns of activity across their voxels. For each parcel and participant, we trained a separate logistic regression classifier using the full multivoxel pattern (*SI Appendix*, Fig. S3*A*). Multivoxel patterns distinguished Chinese from English in 152 parcels (mean accuracy = 54.0%), with consistent decoding across participants distributed across lateral frontal, frontal-opercular/insular, precentral, and medial parietal regions (t(76) ≥ 5.2; Fig. 1*C* ; *SI Appendix*, Fig. S3*B*). At the individual level, significant parcels were sparse and varied substantially across participants (*SI Appendix*, Fig. S4). Searchlight decoding without parcel boundaries found similar regions (*SI Appendix*, Fig. S5).

We used two control analyses to determine how this regional information related to mean parcel activity. First, we reduced each parcel to its mean activity across voxels. The parcel mean rarely distinguished the two languages on its own (19 parcels; mean accuracy = 51.9%; Fig. 1*C*). Second, we removed the parcel mean from the multivoxel pattern before decoding. Chinese and English could still be distinguished in 135 parcels (mean accuracy = 52.4%; Fig. 1*C*). Individual parcels thus carried language state information in fine-grained patterns across their voxels, much of which remained after mean activity was removed, whereas their mean activity rarely distinguished the two languages. Yet the same parcel means, combined into a single cortex-wide pattern, distinguished Chinese from English reliably (59.6%; Fig. 1*B*). Language state is therefore expressed at the scale of the whole cortex, in a pattern that individual parcel means rarely reveal.

Language switching affected the strength of this language state distinction. Chinese and English could be distinguished both when the language repeated from the preceding trial and when it switched, although decoding was stronger on repeat trials (difference 2.8 percentage points, 95% CI [0.6, 5.1]; Fig. 1*B*). Given this effect, we then tested whether switching itself could be decoded from the same naming trials. Yuan et al. [34] previously demonstrated that switch and repeat trials can be distinguished from regional activity patterns. We found no evidence for switch decoding at the cortex-wide scale (49.7%; Fig. 1*B*). Thus, a recent language switch changed how strongly the current language state was expressed without producing a separable cortex-wide switch state.

We used two network definitions to test if language state information was preferentially associated with a particular functional system. First, the Yeo network parcellation [8] provided a broad division of the cortex based on resting-state connectivity. We measured the contribution of each network using permutation feature importance. The parcel features of each network were first reduced with PCA (*Materials and Methods*). For each network, we shuffled its component scores together and measured the resulting drop in decoding performance. A larger drop indicated that the classifier depended more strongly on that network. Contributions were broadly distributed across networks, largest in the somatomotor network (1.9 percentage points [1.3, 2.5]), and lower in the visual network than in the other networks (difference 0.9 percentage points, 95% CI [0.4, 1.4]; Fig. 1*D*; also shown across Yeo 17 networks in *SI Appendix*, Fig. S6). Contributions from the visual and limbic networks did not differ reliably from zero, whereas all other networks contributed reliably (Fig. 1*D*). The finer-grained multivoxel information within individual parcels showed a similar distribution. Informative multivoxel patterns were mostly absent from the visual and limbic networks and were otherwise present in all remaining networks (*SI Appendix*, Fig. S3*C*). Second, we used networks defined from functional localizer experiments in hundreds of participants. This allowed us to compare the language network, which responds selectively during language comprehension, with the multiple-demand network, which is recruited across cognitively demanding tasks [26, 35]. Both have been implicated in bilingual language processing [27, 28, 30]. We tested if the language state classifier depended more strongly on one network than the other. Both contributed reliably to decoding (language: 0.6 percentage points [0.2, 1.0]; multiple demand: 0.7 [0.3, 1.1]), and neither contributed more strongly than the other (difference *−*0.1 percentage points [*−* 0.6, 0.3]; Fig. 1*D*). We found the same balance at the finer regional scale (*SI Appendix*, Fig. S3*C*). The proportion of each network containing parcels that distinguished Chinese from English was similar. For switch status decoding, feature importance across the Yeo networks was centered near zero, with no network showing a clear positive contribution (*SI Appendix*, Fig. S7).

Language state is thus expressed in a cortex-wide pattern of parcel activity, distributed across most networks rather than confined to the language or multiple-demand systems. Decoding accuracy varied widely between participants (43.8% to 73.8%), and informative parcels differed from person to person (*SI Appendix*, Fig. S4). A pattern defined over the same 800 parcels in every brain allows this variation to be examined directly, by testing how well the distinction transfers between people.

### Language state is distinguished along the main axes of task activity

Decoding within a person shows that each brain reliably distinguishes the two language states; transfer between people shows how much of this distinction is shared. We trained a classifier on the full cortical pattern of one participant, the teacher, and applied it to the full cortical pattern of another, the learner. Every participant served in both roles, yielding 5,852 ordered teacher–learner pairs. Language state transferred between people, with a mean accuracy of 56.9% (95% CI [56.0, 57.9]; Fig. 2*B*; *SI Appendix*, Fig. S8), compared with 59.6% [58.4, 60.9] when decoding within the same people across runs (Fig. 3*A*). Part of the neural organization distinguishing the two languages therefore corresponds across bilingual speakers, and part does not.

**Figure 2.**
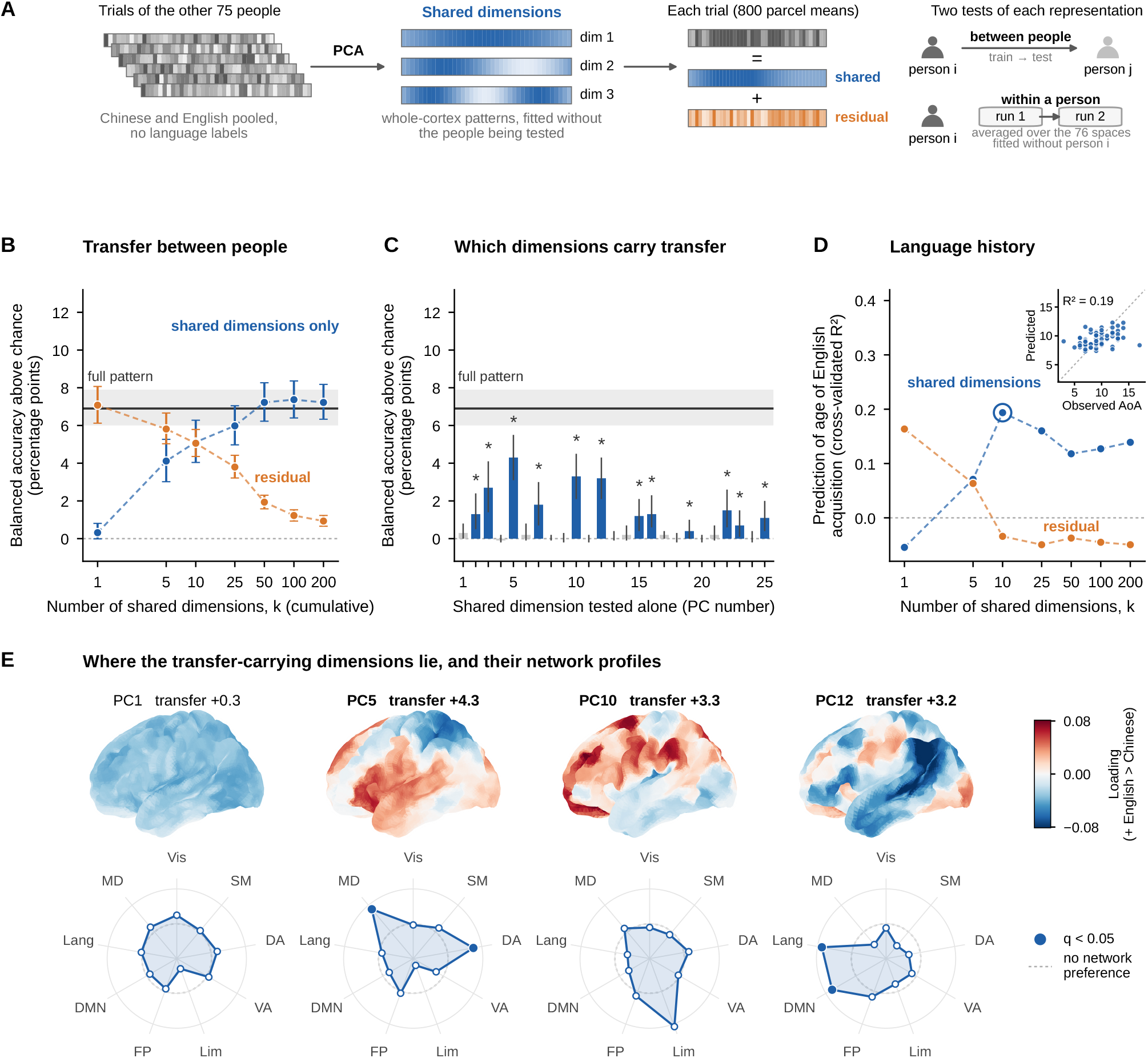
Shared dimensions carry transfer between people, and language history is expressed along them. (*A*) Schematic of the analysis. Shared dimensions are estimated by principal component analysis from the pooled trials of the other 75 participants, without language labels. Each trial is divided into its projection on the first *k* shared dimensions and the residual, and both are tested between people and within each person; within-person results are averaged over the 76 spaces fitted without that person. (*B*) Transfer between people as a function of *k*, using only the first *k* shared dimensions or the residual after removing them. The black line and gray band show transfer with the full cortical pattern; error bars, 95% participant-bootstrap CIs. (*C*) Transfer with each of the first 25 shared dimensions alone; asterisks and blue bars, above chance after Benjamini–Hochberg correction across the 25 dimensions (*q <* 0.05); error bars, uncorrected 95% CIs. (*D*) Prediction of age of English acquisition in held-out participants from learner transfer along the first *k* shared dimensions or the residual. Inset, observed against predicted values at the circled *k* = 10 (*n* = 77). The *k* axes in *B* and *D* are logarithmic. (*E*) Left-hemisphere loading maps of the first dimension and of the three dimensions that carried the most transfer, from a PCA on all participants and signed so that positive values indicate higher scores for English (same color scale for all maps; transfer in percentage points above chance). Radar plots show each network’s share of the dimension’s squared loadings relative to its share of parcels (dashed circle, no network preference, where a network’s share of the dimension equals its share of the cortex; points outside the circle indicate concentration in that network; Vis, visual; SM, somatomotor; DA, dorsal attention; VA, ventral attention; Lim, limbic; FP, frontoparietal; DMN, default; Lang, language; MD, multiple demand). Filled dots mark networks above a spatial null (BrainSMASH, 1,000 surrogates; Benjamini–Hochberg *q <* 0.05 across networks).

**Figure 3.**
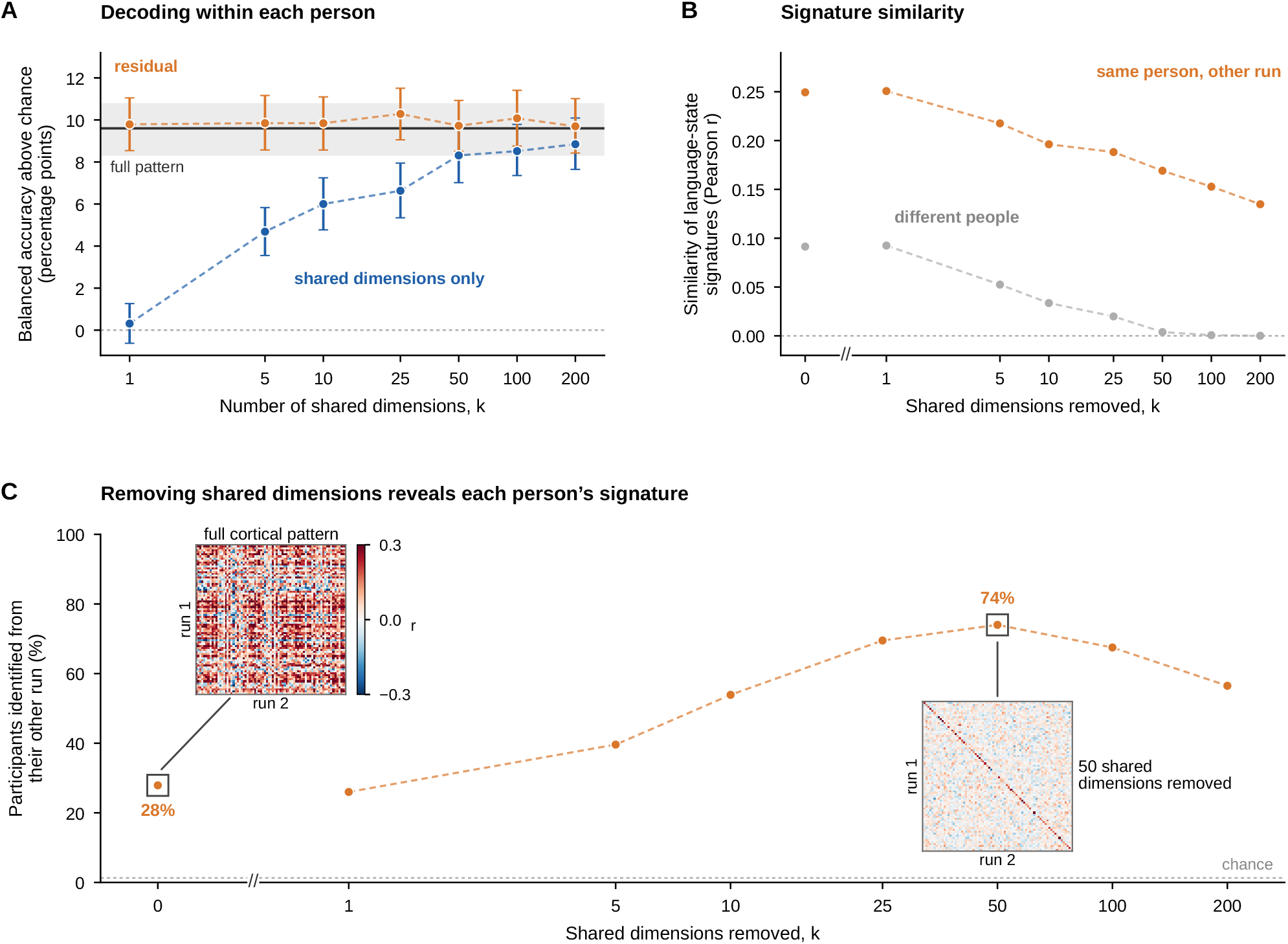
What does not transfer is a stable, person-specific signature. (*A*) Decoding within each person across runs, using only the first *k* shared dimensions or the residual after removing them. The black line and gray band show within-person decoding with the full cortical pattern; error bars, 95% participant-bootstrap CIs (5,000 resamples). (*B*) Similarity (Pearson *r*) of English-minus-Chinese signatures between the two runs of the same participant and between runs of different participants, after removing the first *k* shared dimensions. (*C*) Removing shared dimensions reveals each person’s signature. Percentage of participants identified from their other run, assigning each signature to the most similar signature from the other run and averaging over both directions; dotted line, chance (1/77). Insets show the similarity between the run-1 and run-2 signatures of all pairs of participants at the two boxed points, with the full cortical pattern (28%) and after removing 50 shared dimensions (74%) (same color scale, clipped at ±0.3); each participant’s own match lies on the diagonal. Diagonal entries are averaged over the 76 leave-pair-out spaces that exclude each participant; off-diagonal entries use the space that excludes both. The *k* axes are logarithmic; *k* = 0 (no dimensions removed) is shown separately, after the axis break.

What makes the language-state distinction transferable from one person to another? All participants performed the same task, so the demands it imposes, from recognizing the picture and the cue to selecting the language and producing the word, should shape activity in similar ways across people. For this, we identified the main axes along which task activity varied across participants, without reference to language, and asked whether the distinction between languages lies along them. For every pair of participants, we applied principal component analysis to the pooled naming trials of the remaining 75 participants, combining Chinese and English trials, and projected the pair onto the resulting dimensions. Each of the resulting 2,926 PCA spaces served both transfer directions within its pair, so neither the teacher nor the learner contributed to the dimensions used to test their transfer. We refer to these as *shared dimensions* (Fig. 2*A*). These dimensions show how activity varies around each person’s mean response during the task. We trained classifiers on one person’s Chinese and English trials using only these dimensions, and tested whether they were sufficient to separate the languages in another person.

Transfer required several shared dimensions but far fewer than the full pattern (Fig. 2*B*; *SI Appendix*, Table S1). The first dimension alone carried almost none (50.3% [50.0, 50.8]). Transfer rose with more dimensions (5: 54.1%; 10: 55.1%; 25: 56.0%). The first 10 dimensions captured about half of the variance in task activity (49.0%; *SI Appendix*, Fig. S9). At 50 dimensions it matched and slightly exceeded the full cortical pattern (57.2% [56.2, 58.3]; difference +0.3 percentage points [0.1, 0.5]), and remained there up to 200 (100: 57.4%; 200: 57.2%). Each dimension is a whole-cortex pattern, so this reduction concerns the number of independent patterns needed for transfer, not the extent of cortex involved. The distinction between languages therefore lay along the main axes of task activity, found without language labels, and these axes carried all of the transfer supported by the full cortical pattern. The remaining dimensions could contain only noise, or language information that does not correspond across people. We distinguished these possibilities by removing the shared dimensions.

To identify which shared dimensions carried transfer, we tested each of the first 25 separately (Fig. 2*C*). Twelve transferred above chance after correction, and the largest came from dimensions 5, 10, 12 and 3 (4.3, 3.3, 3.2 and 2.7 percentage points). No single dimension approached the full cortical pattern (6.9), so the distinction between languages was a combination of several axes rather than any single one. To see where these dimensions lie in the cortex, we examined which networks each one drew on. These dimensions had distinct cortical profiles (Fig. 2*E*). PC5, which carried the most transfer, was concentrated in the multiple-demand and dorsal attention networks, and PC12 in the language and default networks (spatial null, corrected). PC1, which carried almost no transfer, loaded with the same sign across the whole cortex, and neither PC1 nor PC10 showed a reliable network preference.

### Age of English acquisition is expressed along the shared dimensions

All participants were non-native speakers of English and Chinese-dominant, and their age of English acquisition (AoA) ranged from 3 to 17 years. We then tried to predict AoA from each participant’s transfer or within-person decoding, testing 43 candidate measures with leave-one-participant-out ridge regression and a max-statistic permutation test (*Materials and Methods*). The best predictor was learner transfer along the first 10 shared dimensions, that is, how accurately classifiers trained on other people decoded a participant’s language state. Participants who acquired English later were decoded more accurately. This measure predicted AoA in held-out participants (*r* = 0.44, cross-validated *R*^2^ = 0.19; Fig. 2*D*) and survived correction across all 43 measures (corrected *p <* 0.001; *SI Appendix*, Table S2). The relationship weakened as more dimensions were included, was absent in the residual (Fig. 2*D*), and was not seen for within-person decoding (*r* = 0.12, cross-validated *R*^2^ = 0.002). Chronological age, which was correlated with AoA, was predicted less well (best cross-validated *R*^2^ = 0.055, corrected *p* = 0.086; *SI Appendix*, Table S3).

### What does not transfer is stable within each person

Removing the shared dimensions reduced transfer progressively (Fig. 2*B*). Removing only the first dimension did not reduce transfer (57.1%). Removing the first 5 lowered it to 55.8%, and removing 50 or 200 brought it close to chance (51.9% and 50.9%; 200 removed: 0.9 percentage points above chance [0.7, 1.2]). The two curves crossed at 10 dimensions, where keeping and removing them supported transfer at a similar level (55.1%).

On the contrary, decoding within individuals stayed just as good after the shared dimensions were removed (Fig. 3*A*). Across runs of the same participant, Chinese and English remained as distinguishable in the residual pattern as in the full cortical pattern at every number of removed dimensions (10 removed: 59.8% [58.6, 61.1]; 200 removed: 59.7% [58.4, 61.0]; full pattern: 59.6%), with no reliable difference at any *k*. The residual therefore still contained the language distinction, but in a form that did not carry over to other people.

If what does not transfer were noise, it could not identify a person. We computed each participant’s language-state signature by subtracting the average brain activity on Chinese trials from that on English trials. We calculated this contrast separately for each run, and assigned each signature from one run to the participant whose signature from the other run correlated with it most strongly, a test used to identify individuals from patterns of brain connectivity [1]. We averaged accuracy over the two directions. With the full cortical pattern, the correct person was identified among 77 in 27.9% of cases (chance, 1.3%). Removing the shared dimensions made individuals easier to identify (Fig. 3*C*). Identification rose as dimensions were removed (10 removed: 53.9%; 25 removed: 69.5%) and was highest after removing 50 (74.0%, 57 of 77), then declined as further dimensions were removed (100: 67.5%; 200: 56.5%). Identification exceeded chance at every step, in both directions (permutation *P <* 0.001).

Identification rose because removing the shared dimensions affected similarity within and between people differently (Fig. 3*B*). Before any dimensions were removed, signatures were more similar within participants (*r* = 0.25) than between participants (*r* = 0.09). As shared dimensions were removed, similarity between participants fell to near zero (50 removed: *r* = 0.004), whereas similarity within participants remained clearly positive (50 removed: *r* = 0.17; 200 removed: *r* = 0.13). How the shared dimensions mask what is individual can be seen more clearly in the matrices of Fig. 3*C*, across all 77 participants. Once 50 shared dimensions were removed, each participant’s match with their own signature became more visible, as can be seen on the diagonal.

## Discussion

Bilingual language use requires selecting one language and producing words in it [16–18]. Every bilingual is unique in the way they have acquired and use their languages [12, 13]. To better understand how the brain distinguishes two languages, we took an approach that identifies consistent patterns in the group and uses them to isolate the remaining part of the signal, in which we find a stable, individual-specific signature.

Here, first of all, we moved beyond regional multivariate pattern analysis to a cortex-wide signal. This choice was well suited to studying individual brain differences, as it avoids selecting regions and missing differences in how language is represented outside the regions picked. In our case especially, the demands associated with English as a second language and the task processes involved in picture naming gave us every reason not to limit our search space. That said, we also reproduced the multivoxel result of previous regional studies [24, 25], and found that parcels distinguished the languages one at a time from their fine-grained voxel patterns (Fig. 1*C*). What was striking is that when we averaged within parcels, discarding fine-grained information [36], parcels rarely had discriminative power on their own. They did so only when combined into a cortex-wide pattern (Fig. 1*B*). Part of this may reflect many weak signals adding up across the cortex. Yet the distinction held across parcellations of different granularity (*SI Appendix*, Fig. S2), which points to a representation of the language in use at another scale.

We found this distinction to be distributed across systems— spanning somatomotor, control, attention, default, language and multiple-demand networks—rather than confined to the language or multiple-demand systems. Given that the language was cued by color, the visual network could have driven the distinction. Our network importance analysis showed that it did not contribute reliably (Fig. 1*D*). This suggests that most of what we capture is not due to the cue color, although cue effects carried by attention or control networks cannot be fully ruled out. Why, then, would the distinction span so many systems? Naming a picture aloud in Chinese or in English requires different articulatory programs—including the laryngeal control of Mandarin tones, which is organized in ventral sensorimotor and laryngeal motor cortex [37–39]—and speaking a second language places additional demands on this system [40, 41]. This may explain why the somatomotor network showed the largest estimated contribution. At the same time, the language network, which supports word meaning and form [26, 27], and the multiple-demand network, which is engaged when a less dominant language must be selected and produced [18, 29, 30, 35], contributed comparably, with neither contributing more than the other. Note that the language state is therefore not exclusively linguistic, it instead appears to be the joint state of several systems.

What role does switching play in this distinction? Chinese and English were distinguished both when the language repeated and when it switched, although less strongly after a switch (Fig. 1*B*). Yet switch and repeat trials could not be told apart at the cortex-wide scale, whereas regional activity patterns have been shown to distinguish them [34]. A recent switch thus weakened how strongly the current language was expressed, without producing a separate cortex-wide state. One possibility is that switching itself is carried by more local signals.

The fact that a classifier trained on one participant could identify the language in another shows a degree of correspondence between people. We show that this correspondence lies along the main axes of task activity (Fig. 2*B*). These axes were estimated without reference to language, but they did indeed provide sufficient correspondence for this transfer. In contrast, the activity remaining beyond these axes transferred close to chance between people, yet it still distinguished the languages within each individual, in a stable and person-specific way (Fig. 3*A* and *C*). This provides an updated framework for interpreting the substantial individual variability reported in bilingual neuroimaging [13–15].

Why should a classifier trained on one bilingual identify the language in another at all, and why only in part? Our results suggest that the distinction is carried by two parts of the same activity pattern. The first lies along the main axes on which all brains vary during the task, which plausibly reflect demands that the task imposes on everyone—recognizing the picture, selecting the language and producing the word. The distinction between languages is a particular combination of these axes, learned from one person and recognized in another. Some of the axes that carried the most transfer were concentrated in systems that also carried the distinction within each person, one in the multiple-demand and dorsal attention networks and another in the language and default networks. Others showed no clear network preference, and none was concentrated in the somatomotor network (Fig. 2*E*). The remaining part belongs to each person and may reflect how each meets the same demands, including individual strategies for selecting and producing words in each language as well as individual differences in functional organization. Although it transfers poorly, it distinguishes the languages within that person and recurs across runs. It is important to note, therefore, that lower accuracy across people than within a person does not mean that the distinction is weak in individual brains; rather, it reflects how much of each person’s distinction lies on the shared dimensions. Aligning fine-grained patterns between people has revealed shared representations for perceived content such as familiar faces [42]. Here, a shared representation exists for a state the speaker produces, at a scale where anatomy alone is enough to compare brains, unlike fine-grained information, which requires functional alignment [43, 44]. Similarly, in the brain at rest, removing what is common to all people makes individual differences in how regions fluctuate together easier to detect [45]. We find that the same holds during a task: removing the shared dimensions raised the identification of individuals from their language-state signature from 28% to 74% (Fig. 3*C*). Identification declined again when more than 50 dimensions were removed. Part of what is individual may therefore also lie on the higher shared dimensions.

A person’s language history, indexed here by the age of English acquisition, gave insight into what the shared axes might reflect. Although modest (cross-validated *R*^2^ = 0.19), it explained part of why some people transferred better than others along the shared dimensions (Fig. 2*D*). Later learners of English were decoded more accurately by other people’s classifiers, although their own decoding across runs was not stronger. A later-acquired second language is known to rely more on control processes in prefrontal cortex [18, 29, 30]. The extent of this reliance may lie along a continuum captured by the shared axes, such as PC5, which was concentrated in the multiple-demand and dorsal attention networks (Fig. 2*E*). Age of acquisition was not reflected in the person-specific residual, suggesting that it particularly links to the shared organization we find here.

These results were obtained with simple linear methods and a limited amount of data, two runs and 160 trials per person. Studies of individual brains have often collected many hours of data from each person [2, 3, 11], which many fields, including bilingualism, cannot do. Decoding a cognitive state in a common anatomical frame, and separating what transfers from what stays within a person, offers a way to study shared and individual organization with the data such fields typically have.

These results should be interpreted in light of the design of the study. Naming responses were not recorded in the scanner, although accuracy on the same task outside the scanner was high [31]. The task involved naming 40 pictures in rapidly alternating languages. Overlapping responses between neighboring trials may therefore have weakened the distinction, and slower designs could reveal the distinction more fully. Our analyses were also restricted to the cortex. Does the cerebellum, which also contributes to bilingual language control [46], carry part of the distinction? We don’t yet know. More generally, the results come from one dataset and one language pair. One difficulty here is that other available datasets present different languages to different groups of participants [47] or in separate blocks [30], which does not allow the trial-level decoding used here. Whether the same shared and person-specific organization holds for other language pairs, tasks and cognitive distinctions would require additional investigation.

To conclude, every bilingual must select one language and produce words in it, and each has acquired and uses their languages in their own way [12, 13]. Here, we decoded the language in use from cortex-wide patterns of activity and showed that the distinction between a bilingual’s languages is expressed at the scale of the whole cortex. Parcel means that rarely distinguished the languages one at a time did so reliably when combined into a cortex-wide pattern. This distinction was partly shared across individuals, as a classifier trained on one bilingual identified the language in use in another. The shared part lay along the main axes of task activity common to all participants, and age of English acquisition was expressed along these axes. The activity remaining beyond this shared part was not simply noise but a stable signature specific to each person. This work suggests that how the brain distinguishes a bilingual’s languages is partly common to all speakers and partly their own. By separating what transfers between people from what stays within each person, future work should continue to examine how shared and individual organization support other languages, tasks and cognitive states.

## Materials and Methods

### Dataset

We analyzed data from 77 Chinese–English bilinguals (34 female, 43 male; mean age = 22.2 years, standard deviation = 2.2) from the dataset of Guo et al. [31], publicly available on OpenNeuro (ds005455, version 1.1.5). Participants formed a relatively homogeneous group in their language acquisition, proficiency, and use. They were Chinese-dominant bilinguals living in China who began learning English at around nine years of age, primarily through school. All participants provided written informed consent to participate in the study and to publicly share their de-identified data. The original study was approved by the Institutional Review Board of the Imaging Center for Brain Research at Beijing Normal University (Protocol ICBIR A 0012 003). More details on recruitment, inclusion criteria, and data acquisition are provided with the original dataset.

### Neuroimaging task

Participants completed a cued picture-naming task during functional magnetic resonance imaging (fMRI) [31]. During the task, participants named a line drawing in either Chinese, their dominant and first language (L1), or English, their less dominant and second language (L2). A red or blue frame around the picture indicated which language to use, with color assignments counterbalanced across participants. The task required participants to repeatedly alternate between Chinese and English. Each picture was presented for one second, followed by an interstimulus interval of one to four seconds. The task design is illustrated in *SI Appendix*, Fig. S10. Each naming trial was labeled by the cued language and switch status. L1S and L2S (S for “switch”) mean that the cued language differed from that of the preceding trial. L1NS and L2NS (NS for “non-switch”) mean that the language remained the same as on the preceding trial. Participants named each picture aloud during scanning, but verbal responses were not recorded in the scanner for technical reasons [31]. Behavioral performance was assessed outside the scanner, where participants repeated the task and their naming responses were recorded (*SI Appendix*, Fig. S11).

### Data preprocessing

We used the single-trial blood-oxygen-level-dependent (BOLD) activation estimates released with the dataset of Guo et al. [31]. These images were generated by the dataset authors using a Least Squares-Separate (LSS) general linear model, resulting in one trial-specific activation estimate for each picture-naming trial. Details of MRI acquisition, image preprocessing, and single-trial model estimation are reported in the original dataset paper. For each participant, we used the single-trial images from the two LanguageControl runs together with their corresponding event files. Images were resampled to the 2-mm Montreal Neurological Institute (MNI152) template used in the present analyses when voxelwise or parcelwise feature extraction required a common grid.

### Cortex-wide decoding

Cortex-wide decoding represented each trial by the pattern of mean activity across Schaefer parcels. For the main analysis, we used the Schaefer-800 parcellation [9], giving one feature per parcel. The dataset contained single-trial blood-oxygen-level-dependent (BOLD) activity estimates for 160 naming trials across two experimental runs. The single-trial images were in Montreal Neurological Institute (MNI) space but had a different voxel grid from the 2-mm Schaefer atlas, so the atlas labels were resampled to the single-trial image grid during feature extraction. For each participant, the single-trial images were concatenated and passed to NiftiLabelsMasker, without standardization. BOLD activity was averaged across voxels within each parcel, so each trial was represented by 800 parcel-mean activity values. Neuroimaging feature extraction and visualization were performed using Nilearn version 0.13.1 [48]. Chinese picture-naming trials (L1S+L1NS) were classified against English picture-naming trials (L2S+L2NS) using L2-regularized logistic regression (C=0.01, solver=lbfgs, max iter=1000). Decoding performance was measured using balanced accuracy. We used leave-one-run-out cross-validation, training the classifier on trials from one experimental run and testing it on trials from the other. Features were z-scored within each cross-validation fold using parameters estimated from the training run only. Chance was 50%. Group means are reported with 95% confidence intervals obtained by bootstrapping participants [49]. We drew 77 participants at random from the sample 5,000 times, allowing the same participant to be drawn more than once. We recomputed the mean at each iteration. The interval is the 2.5th to 97.5th percentile of these means. We assessed differences between conditions and differences from chance by bootstrapping the paired differences. A difference was considered reliable when its 95% confidence interval excluded zero. The same procedure was used for all confidence intervals and comparisons below. The relationship between decoding accuracy and mean temporal signal-to-noise ratio across participants was assessed with Pearson and Spearman correlations. *Robustness to cortical representation*. We tested if cortex-wide decoding depended on how cortical activity was represented. First, we varied the spatial granularity of the parcellation by repeating the analysis with Schaefer atlases ranging from 100 to 1000 parcels. At each granularity, a trial was represented by the pattern of mean activity across parcels. Second, we tested the effect of dimensionality reduction using principal component analysis (PCA). PCA was fit to the training data within each cross-validation fold, and components explaining 90% of the training-set variance were retained. The training and test data were projected onto these components, and classification was performed using the resulting component scores. *Decoding across switching contexts*. We tested if the distinction between Chinese and English was present both when participants continued using the same language and when they had just switched languages. Chinese versus English decoding was repeated separately for switch trials (L1S versus L2S) and nonswitch trials (L1NS versus L2NS). Each analysis contained 80 trials instead of the 160 trials used in the main analysis. Decoding on repeat and switch trials was compared with the bootstrap on the paired difference. *Switch-status decoding*. We separately tested if the same cortex-wide activity patterns distinguished switching itself. Trials were relabeled according to switch status, and classifiers distinguished switch trials (L1S + L2S) from nonswitch trials (L1NS + L2NS). Classification otherwise followed the same cortex-wide decoding procedure.

### Network importance

We tested if particular functional systems contributed more strongly to cortex-wide language state decoding. We used two complementary network definitions. The Yeo networks provide a broad division of the cortex based on resting-state functional organization [8]. We used the Yeo 7-network parcellation for the main analysis. We examined the Yeo 17-network parcellation as a supplementary analysis. The networks defined by Yeo parcellation were non-overlapping, so each parcel belonged to a single network. For a functional comparison, we used the probabilistic language network atlas [26] and multiple-demand network atlas [35], obtained from the EvLab resources repository. We resampled both maps to Schaefer-800 space and averaged the probabilities within each parcel. For each atlas, parcels were ranked by their mean probability, and the top 10% were retained (80 parcels per set). The resulting Lipkin Language and Lipkin Multiple-Demand sets did not overlap with each other. These sets were defined separately from the Yeo networks, so a parcel could belong to both a Yeo network and either the Lipkin Language or Lipkin Multiple-Demand set. The spatial definitions of all network and functional parcel sets used in these analyses are shown in *SI Appendix*, Fig. S12. We quantified the contribution of each network to whole-cortex decoding using permutation feature importance. Within each cross-validation fold, the parcel features of each network were z-scored and reduced with PCA, keeping the components that explained 90% of that network’s variance. Both steps were fitted to the training run only, and the held-out run was projected onto the resulting components. The component scores of all networks were concatenated, and a single classifier was trained on them. For each network, we shuffled its component scores together across held-out trials 20 times and recomputed decoding accuracy using the same trained classifier. Network importance was defined as the mean decrease in accuracy caused by this shuffle, averaged across shuffle repeats and cross-validation folds within each participant. Larger decreases therefore indicated stronger dependence on information carried by that network (Fig. 1*D*; *SI Appendix*, Figs. S6*B* and S7*B*). *Size-matched importance*. As a complementary estimate, we also computed network importance directly on the parcel features, without PCA (*SI Appendix*, Figs. S6*A* and S7*A*). To make network importance comparable across the Yeo networks, we matched them to the size of the smallest network. The smallest Yeo network, Limbic, contained 54 parcels. We therefore compared networks using the same number of parcels, repeatedly sampling 54 parcels from each larger network 500 times. For each cross-validation fold and each size-matched sample, activity from the selected parcels was shuffled across held-out trials 20 times. Consequently, network importance was determined based on 20,000 permutations for each larger network and each participant. Permutation importance was defined as the mean decrease in decoding accuracy relative to the unshuffled held-out accuracy. Importance values were averaged across shuffle repeats, size-matched samples, and cross-validation folds within each participant. For both estimates, differences between networks were assessed on these participant-level importance values with the bootstrap described above, comparing the visual network with the mean of the other Yeo networks, and the language set with the multiple-demand set. In *SI Appendix*, Figs. S6 and S7, each network was compared with the visual network in the same way.

### Multivoxel decoding

We tested if Chinese and English could also be distinguished from finer-grained patterns of voxel activity within cortical parcels. We used L2-regularized logistic regression (lbfgs, C = 0.01, max iter = 1000) and measured decoding performance using balanced accuracy. Decoding used leave-one-run-out cross-validation: the classifier was trained on trials from one experimental run and tested on trials from the other. Features were z-scored within each cross-validation fold using parameters estimated from the training run only. *Parcelwise multivoxel decoding*. For each participant and each of the 800 Schaefer parcels, the BOLD activity of each voxel was entered as a separate feature, allowing the classifier to use the full pattern of voxel activity within the parcel. Group-level inference tested parcelwise balanced accuracies against chance (50%) using one-sided one-sample t-tests across participants, followed by Benjamini–Hochberg false discovery rate (BH-FDR) correction [50] across the 800 parcels (q ≤ 0.001). Participant-level significance was assessed using one-sided binomial tests on held-out predictions, followed by BH-FDR correction across the 800 parcels within each participant (q *<* 0.05) (*SI Appendix*, Fig. S4). *Control analyses of parcel representation*. We repeated parcelwise decoding using two alternative representations. To test if the fine-grained voxel pattern remained informative after differences in mean parcel activity were removed, we represented each parcel by its mean activity across voxels and performed decoding using a single feature per trial. For our second control, we removed the mean activity of the parcel from every voxel on each trial and repeated decoding using the resulting demeaned multivoxel pattern. Classification, cross-validation, and statistical inference were kept identical to the main parcelwise analysis.

### Cross-participant transfer and shared dimensions

*Cross-participant transfer*. For each ordered pair of participants, a classifier was trained on all trials of one participant (the teacher) and tested on all trials of the other (the learner). In total, we had 77 *×* 76 = 5,852 transfer models, both learner-teacher directions combined. We trained and tested transfer models using three representations of each trial: the full cortical pattern (800 parcel means), the scores on the first *k* shared dimensions, and the residual pattern after removing those dimensions. Each participant’s activity was z-scored per parcel across all of that participant’s trials. All three representations were derived from these z-scored data without further standardization. Classification used the same L2-regularized logistic regression as within-participant decoding. For each participant, learner transfer was the mean accuracy of the 76 classifiers trained on the other participants and tested on this participant. Teacher transfer was the mean accuracy of this participant’s classifier tested on the 76 other participants. Confidence intervals for transfer used the participant bootstrap described above, with the mean of each resample computed over the teacher–learner pairs formed by the resampled participants. Transfer with each of the first 25 shared dimensions alone was tested with the same bootstrap; one-sided *P* values were the proportion of bootstrap means at or below chance and were corrected across the 25 dimensions with the Benjamini–Hochberg procedure [50]. *Shared dimensions*. We identified the dominant dimensions of task activity with principal component analysis (PCA). Given a set of trials, each described by its 800 parcel values, PCA finds a series of whole-cortex patterns, the principal components, each a weighted combination of all 800 parcels. The first component is the pattern along which activity varies most across trials. Each subsequent component captures the most remaining variance while being orthogonal to the previous ones. The first *k* components therefore give the *k*-dimensional description of the activity that preserves the most variance. PCA was fitted to these z-scored data separately for every unordered pair of participants, always leaving out both members of the pair. This gave 77 *×* 76 / 2 = 2,926 PCA spaces, each fitted to the pooled trials of the remaining 75 participants (12,000 trials *×* 800 parcels). Chinese and English trials were pooled, and PCA was not told which language each trial belonged to. We refer to the leading components as shared dimensions. For display and network profiles, PCA was also fitted to the pooled trials of all 77 participants; its first 10 components matched those of every leave-pair-out space (cosine similarity ≥ 0.989). The sign of each component was set so that positive values indicated higher scores for English than for Chinese trials. For each component and network (the seven Yeo networks and the language and multiple-demand sets), enrichment was the network’s share of the component’s squared loadings divided by its share of parcels. Enrichment was compared with 1,000 surrogate maps that preserved the spatial autocorrelation of each component, generated separately for each hemisphere from Euclidean distances between parcel centroids (BrainSMASH [51]), with Benjamini–Hochberg correction across networks within each component. PCA was implemented in scikit-learn 1.8.0 [52] using randomized singular value decomposition with a fixed random seed (random_state = 42). *Transfer in the shared dimensions*. Each trial was represented by its scores on the first *k* shared dimensions, that is, by how strongly each of the *k* whole-cortex patterns was expressed in the trial’s activity. In this case, the classifier received *k* values per trial as input rather than 800 parcel values. For *k* = 1, 5, 10, 25, 50, 100 and 200, it was trained on the teacher’s scores and tested on the learner’s scores. Transfer with the full cortical pattern is mathematically equivalent to transfer with all 800 components, so the full pattern is the endpoint of the same analysis. We verified this numerically for 40 directed pairs, in which transfer with all 800 components and with the full cortical pattern differed by at most one trial. *Transfer in the residual pattern*. Each trial was reconstructed from its scores on the first *k* shared dimensions, and this reconstruction was subtracted from the trial’s activity:

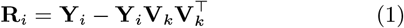

where **Y**_*i*_ is the z-scored trial *×* parcel activity matrix of participant *i* and **V**_*k*_ contains the first *k* shared dimensions as columns (800 *× k*). **Y**_*i*_**V**_*k*_ gives the trial scores, and multiplying by the transpose of **V**_*k*_ rebuilds the corresponding 800-parcel activity. The residual **R**_*i*_ therefore stays in the original 800-parcel space but contains only activity that the first *k* shared dimensions do not capture. The classifier was trained on the teacher’s residual patterns and tested on the learner’s residual patterns, for the same values of *k. Within-participant decoding with shared dimensions*. To compare transfer with decoding inside a single brain, we repeated within-participant decoding (leave-one-run-out, as above) on the full cortical pattern and on the same two representations: scores on the first *k* shared dimensions, and residual patterns after their removal. For this comparison, decoding used the same per-participant z-scored data as transfer, without the fold-wise standardization of the main within-participant analysis. Each participant was excluded from 76 PCA spaces, one for each possible partner. Within-participant decoding was repeated in each of these spaces, and the accuracies were averaged, giving one within-participant accuracy per participant for each representation and *k*. These averaged accuracies are the within-participant measures shown in Fig. 3*A*. *Similarity of residual language-state signatures.* We tested whether the language-state information that remained after removing the shared dimensions was specific to each individual. For each participant and run, we computed a language-state signature: the mean activity pattern across the 800 parcels for English trials minus the mean pattern for Chinese trials. We then correlated (Pearson *r*) each participant’s run-1 signature with every participant’s run-2 signature, giving a 77 *×* 77 matrix of correlations. Diagonal entries, the two runs of the same participant (77 values), gave within-participant similarity; off-diagonal entries, run 1 of one participant with run 2 of another (5,852 values), gave between-participant similarity. Each was averaged across its entries. For identification, each participant’s signature from one run was assigned to the participant whose signature from the other run correlated with it most strongly, and accuracy was the proportion of participants assigned to themselves (chance, 1/77). Identification was computed in both directions and assessed by permuting participant identities across runs (10,000 permutations). The analysis was performed on the full cortical pattern, with no dimensions removed, and repeated after removing the first *k* shared dimensions from each trial as in Eq. 1, for the same values of *k*. Removal used the leave-pair-out PCA spaces, so the participants being compared never contributed to the dimensions removed: between-participant correlations used the space fitted without the two participants compared, and within-participant correlations were averaged over the 76 spaces excluding that participant, as for within-participant decoding.

### Prediction of age of English acquisition

We tested whether individual differences in transfer and in within-participant decoding predicted age of English acquisition (AoA). Three measures were used for each participant: learner transfer and teacher transfer (defined above), and within-participant decoding across runs (see Within-participant decoding with shared dimensions). Each measure was computed on the scores of the first *k* shared dimensions and on the residual patterns after their removal, for *k* = 1, 5, 10, 25, 50, 100 and 200 (42 predictors). Within-participant decoding on the full cortical pattern was included as a 43rd predictor. AoA was predicted separately from each predictor using leave-one-participant-out cross-validation. On each fold, the predictor was standardized using the training participants only, a ridge regression model (*α* = 1) was fitted to the remaining participants, and AoA was predicted for the held-out participant. Prediction performance was summarized by the cross-validated *R*^2^ and by the Pearson correlation between observed and predicted AoA. *Max-statistic permutation test*. To account for testing all 43 candidate predictors, AoA values were permuted across participants 10,000 times [53]. For each permutation, the complete leave-one-participant-out procedure was repeated for every candidate predictor, and the largest cross-validated *R*^2^ was retained. The corrected p value was the proportion of permutations whose maximum *R*^2^ was greater than or equal to the best observed *R*^2^. The same procedure was used with chronological age as the outcome.

## Ethics statement

This study did not collect new data from human participants and analyzed a publicly available, de-identified dataset. In the original study, all participants provided written informed consent to participate and to publicly share their de-identified data, and the study was approved by the Institutional Review Board of the Imaging Center for Brain Research at Beijing Normal University (Protocol ICBIR A 0012 003). No additional institutional ethics approval or consent was required for the present secondary analysis.

## Generative AI statement

Generative AI was used only for editing the text. The authors take full responsibility for the content.

## Data, Materials, and Software Availability

The data are available on OpenNeuro (ds005455). All analysis code is available at https://github.com/Sinem-demirkan/multiscale_bilingual_decoding.

## Acknowledgments

This work was supported by B.S.’s chair in PRAIRIE-PSAI, funded by the French national agency ANR as part of the “France 2030” strategy (ANR-23-IACL-0008).

## Author contributions

S.D., D.W., and B.S. designed research; S.D. performed research; S.D. analyzed data; S.D. wrote the paper; D.W. and B.S. reviewed and edited the paper; and B.S. acquired funding.

