## Supplementary Material for "How the brain distinguishes a bilingual’s languages is partly shared across people and partly their own"

### Supplementary Methods

#### Single-trial data

In the released single-trial derivatives [1], each 4D NIfTI file corresponds to one experimental run, and each volume corresponds to one trial in chronological order.

#### Trial definitions

Each picture-naming trial belonged to one of four experimental conditions defined by the target language and whether the language repeated or switched relative to the preceding trial: L1NS, Chinese nonswitch; L1S, Chinese switch; L2NS, English nonswitch; and L2S, English switch. Here, L1 and L2 denote the first and second language, respectively, whereas NS and S denote nonswitch and switch.

#### Searchlight multivoxel decoding

We also tested for language information without restricting voxel patterns to parcel boundaries. For each participant, the BOLD activity values of all voxels within a moving 4-mm-radius searchlight sphere were entered as classifier features. Classification used the same L2-regularized logistic regression and leave-one-run-out cross-validation procedure as the parcelwise analysis, with balanced accuracy as the performance measure. For group-level inference, participant searchlight accuracy maps were spatially smoothed with a 6-mm full-width-at-half-maximum Gaussian kernel. At each voxel, balanced accuracy was tested against chance (50%) across participants using one-sided one-sample t-tests, followed by BH-FDR correction across voxels ($q \leq 0.001$) (Fig. S5).

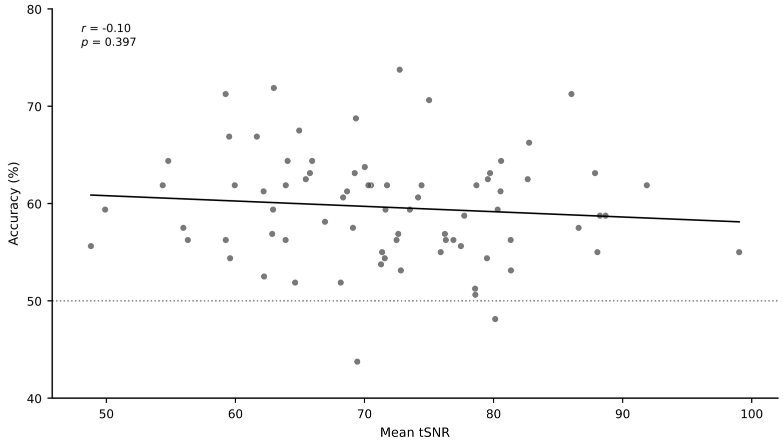


**Figure S1. Whole-cortex decoding accuracy does not depend on signal quality.** Each dot is one participant ($n$ = 77). Within-participant decoding accuracy (leave-one-run-out, Schaefer-800 parcel means) is plotted against mean temporal signal-to-noise ratio (tSNR). The line shows the least-squares fit and the dotted line chance (50%). Pearson $r = -0.10$, $p = 0.40$; Spearman $\rho = -0.10$, $p = 0.38$.

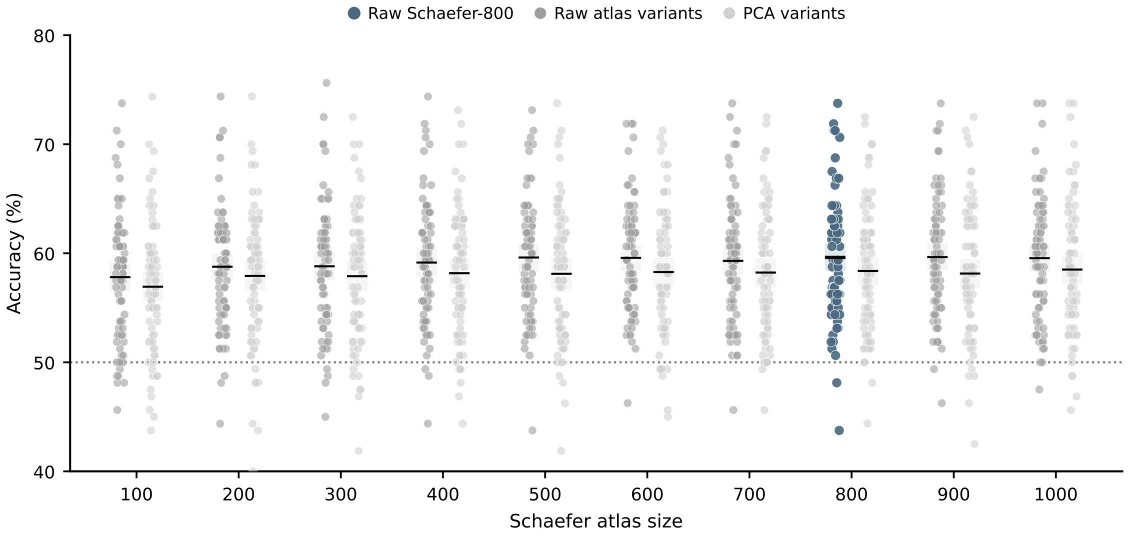


**Figure S2. Decoding is robust to the granularity of the parcellation and to dimensionality reduction.** Within-participant decoding accuracy for Schaefer atlases with 100 to 1,000 parcels, using the parcel means directly (raw) or after PCA retaining the components that explained 90% of the training-set variance within each cross-validation fold (PCA variants). Dots are participants; the main analysis (raw Schaefer-800) is highlighted. Horizontal bars show group means and shaded boxes their 95% bootstrap confidence intervals. The dotted line marks chance (50%).

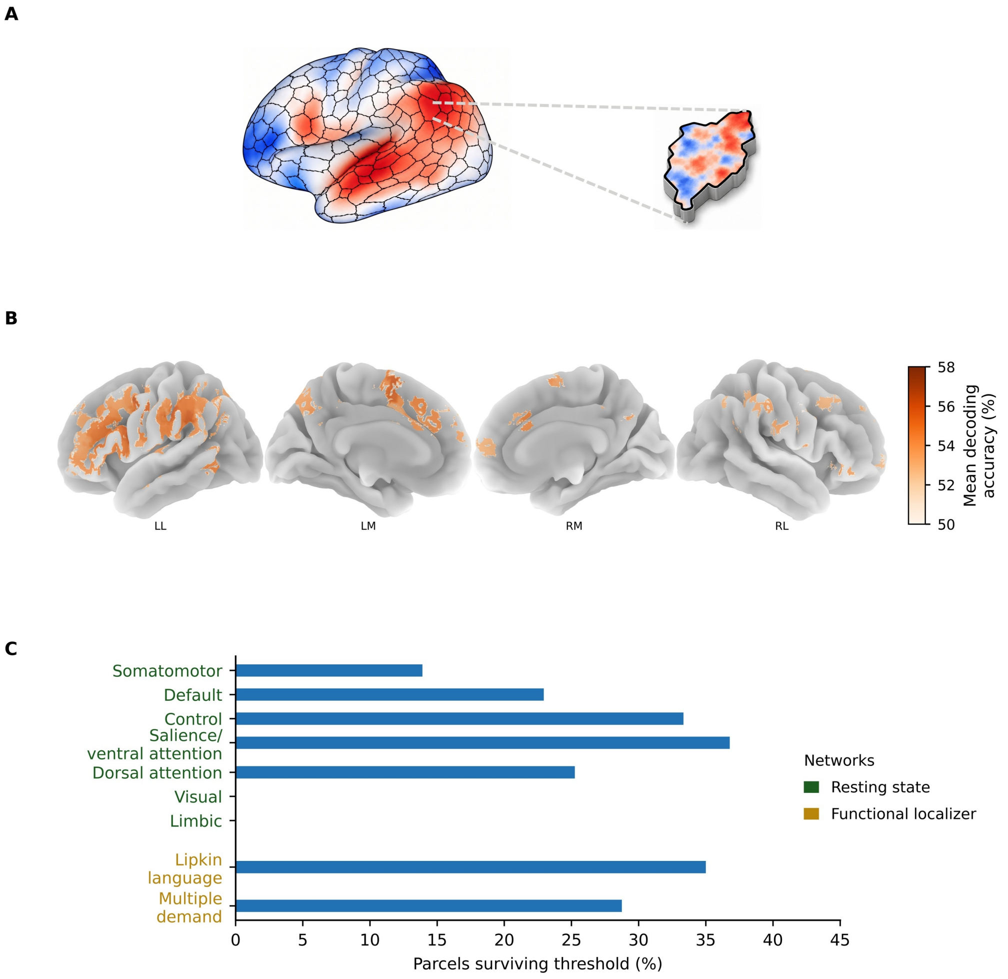


**Figure S3. Multivoxel activity patterns within cortical parcels distinguish between language states.** (*A*) Analysis schematic. Language state decoding was performed separately within each Schaefer parcel using logistic regression. (*B*) Group-level multivoxel decoding map. Surface maps show mean decoding accuracy for parcels surviving the group-level threshold, $t(76) \geq 5.2$. Surface views are left lateral (LL), left medial (LM), right medial (RM), and right lateral (RL). (*C*) Network distribution of multivoxel decoding. Bars show the percentage of parcels that survived the threshold within each of the seven Yeo networks and the functional localizer networks. The two network definitions were analyzed separately and may overlap spatially.

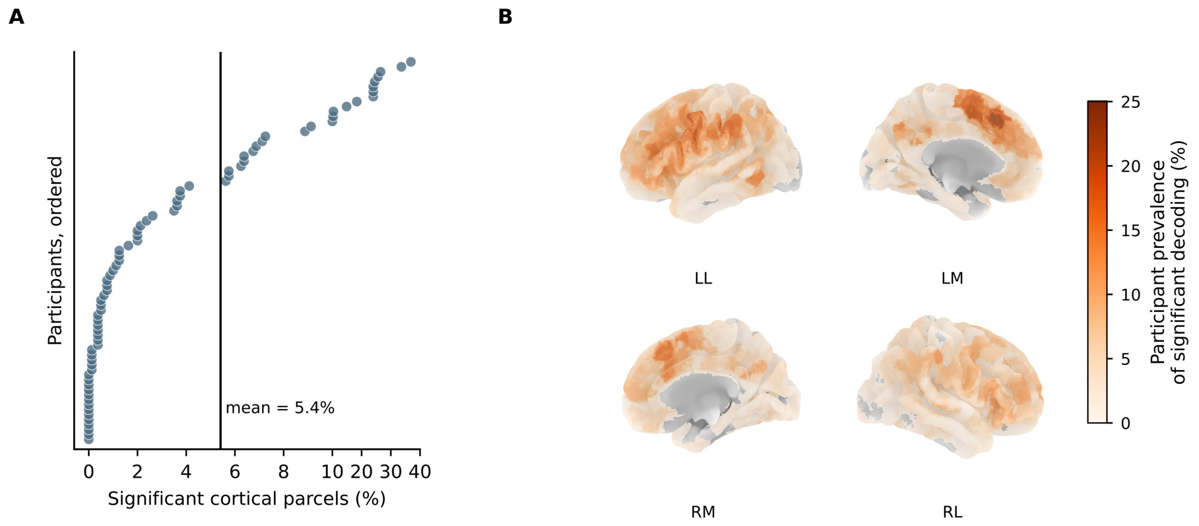


**Figure S4. Informative parcels differ between individuals.** (*A*) Percentage of the 800 parcels whose multivoxel pattern distinguished Chinese from English in each participant (one-sided binomial test on held-out predictions, BH-FDR correction across parcels within each participant, $q < 0.05$), with participants ordered by this percentage. The vertical line marks the mean (5.4%). The horizontal axis is expanded between 0 and 10%. (*B*) Percentage of participants in whom each parcel was significant. Surface views are left lateral (LL), left medial (LM), right medial (RM), and right lateral (RL).

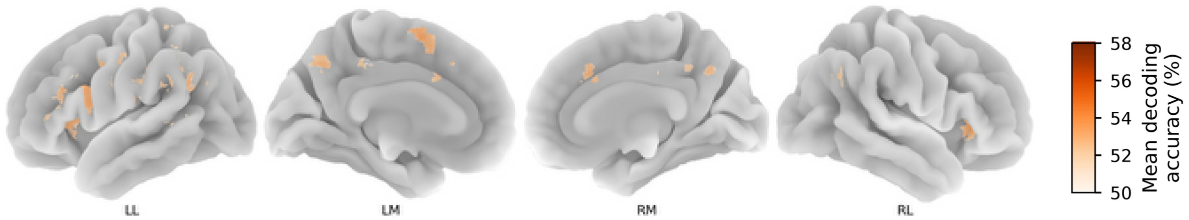


**Figure S5. Searchlight multivoxel decoding.** Group mean decoding accuracy of a 4-mm-radius searchlight, shown at voxels where accuracy exceeded chance across participants (one-sided one-sample $t$ test, BH-FDR $q \leq 0.001$) after smoothing individual accuracy maps with a 6-mm FWHM Gaussian kernel. Surface views are left lateral (LL), left medial (LM), right medial (RM), and right lateral (RL).

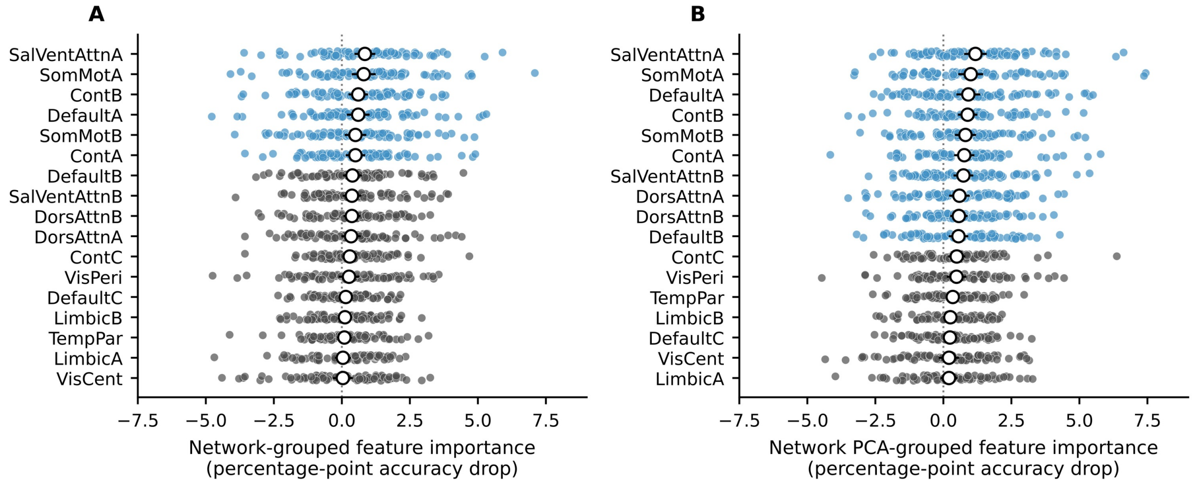


**Figure S6. Network contributions across the Yeo 17 networks.** Permutation feature importance, the drop in decoding accuracy when a network's features were shuffled across held-out trials. (*A*) Size-matched importance, computed directly on the parcel features of each network, with networks matched in size. (*B*) Importance computed on the PCA component scores of each network, as in the main analysis (Fig. 1*D*). Both estimates are described in *Materials and Methods.* Dots are participants; white circles show group means with 95% bootstrap confidence intervals. Blue indicates networks with greater importance than the central visual network (VisCent).

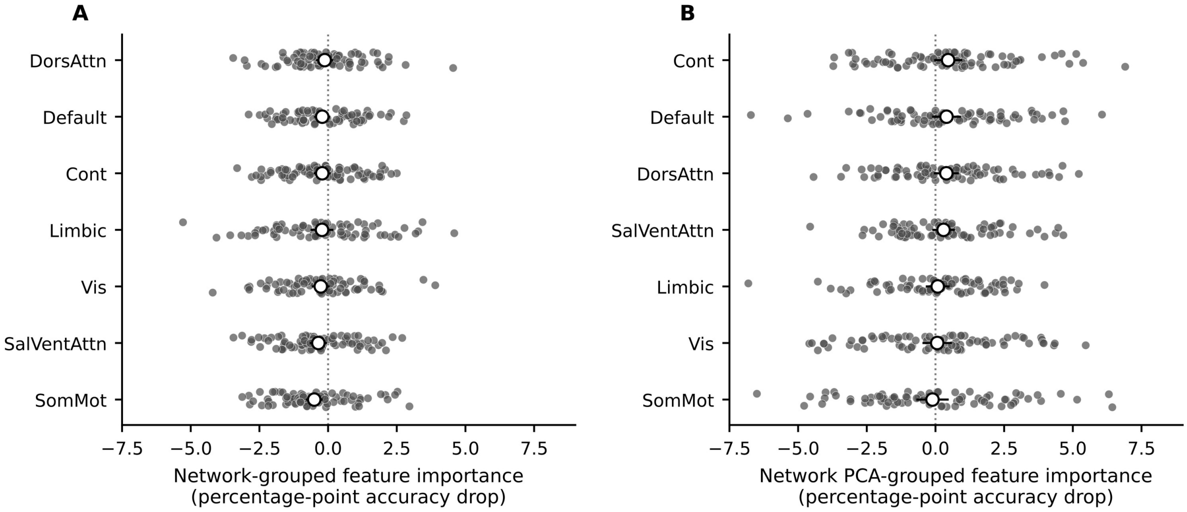


**Figure S7. No network contributes to decoding switch status.** Permutation feature importance for classifiers distinguishing switch from nonswitch trials, shown for the Yeo 7 networks. (*A*) Size-matched importance, computed directly on the parcel features of each network, with networks matched in size. (*B*) Importance computed on the PCA component scores of each network, as in the main analysis. Both estimates are described in *Materials and Methods*. Dots are participants; white circles show group means with 95% bootstrap confidence intervals. Networks with greater importance than the visual network would be shown in blue; none were.

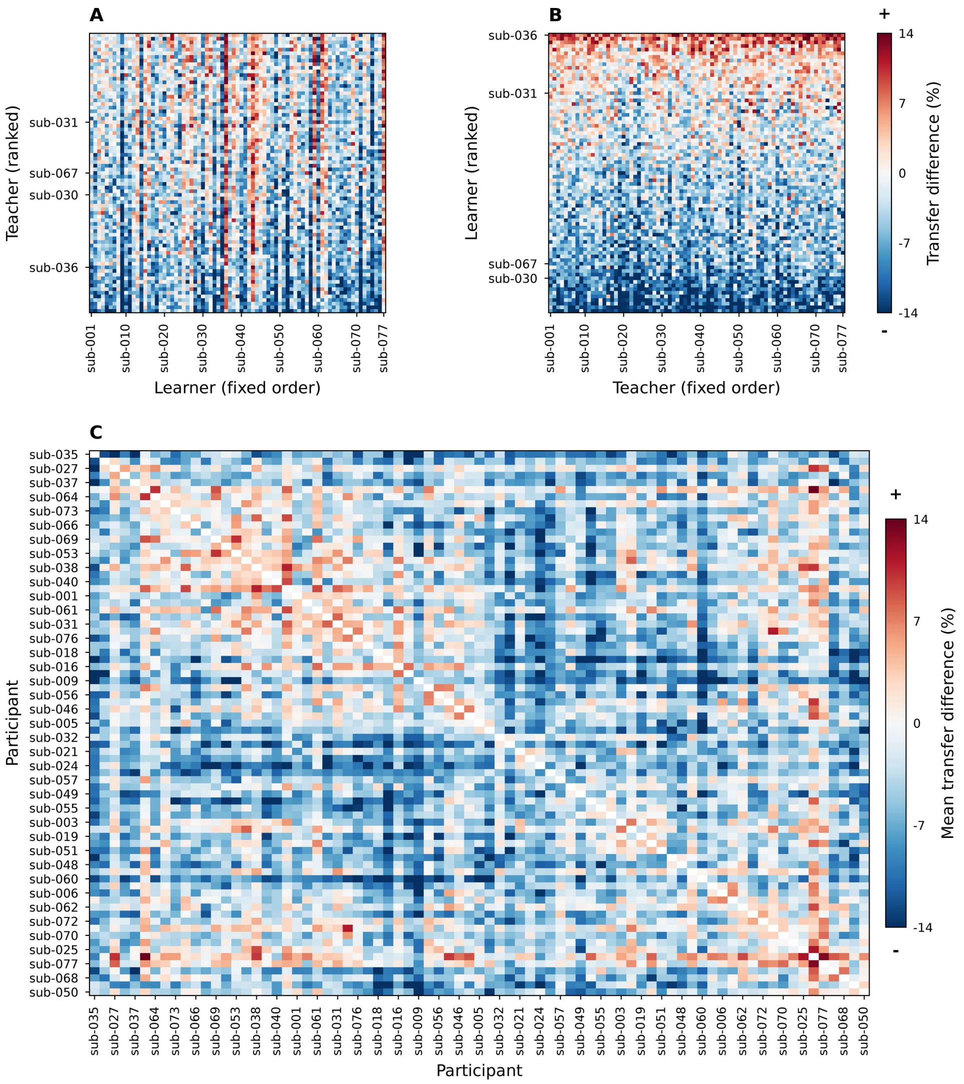


**Figure S8. Transfer between individual pairs of participants.** Each cell shows the transfer difference for one pair: the accuracy of a classifier trained on the teacher and tested on the learner, minus the learner's own within-participant accuracy. Red cells indicate transfer above, and blue cells below, the learner's own decoding. (*A*) Teachers ranked (rows) against learners in a fixed order (columns). (*B*) Learners ranked (rows) against teachers in a fixed order (columns). (*C*) Mean transfer difference for each pair of participants, averaged over the two directions. Labeled participants in *A* and *B* mark the same individuals in both rankings.

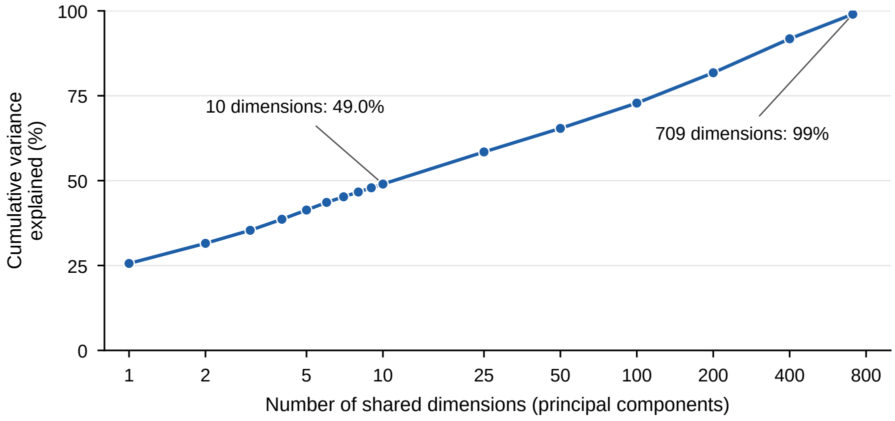


**Figure S9. Variance of task activity captured by the shared dimensions.** Cumulative percentage of variance in the z-scored single-trial activity of all participants explained by the leading principal components (logarithmic horizontal axis). The first 10 dimensions captured 49.0% of the variance, and 709 dimensions were needed to reach 99%.

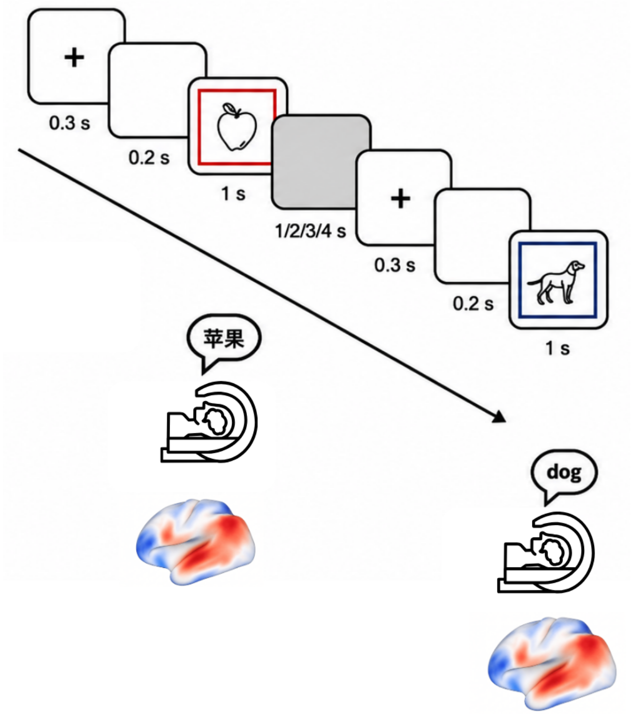


**Figure S10. Neuroimaging picture-naming task.** On each trial, participants viewed an object picture surrounded by a blue or red frame and named the object aloud in Chinese or English according to the frame color. The required language either repeated from the previous trial or switched. Figure adapted from Guo et al. [1], published under a CC BY 4.0 license.

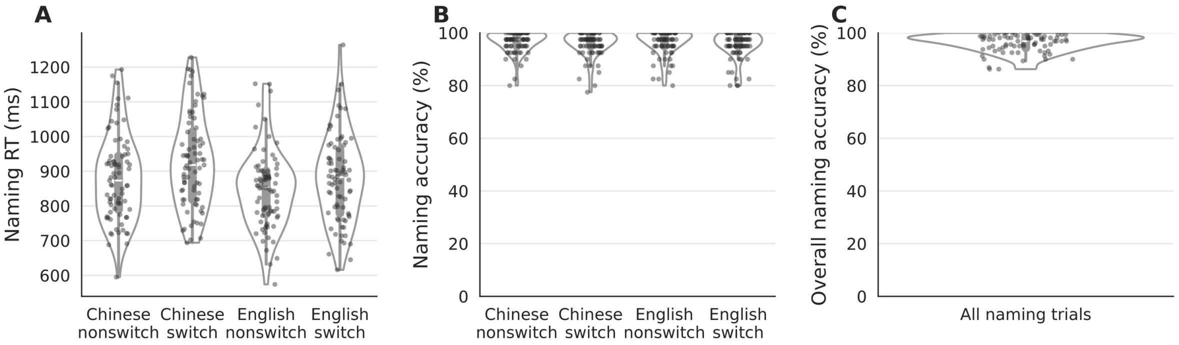


**Figure S11. Naming performance outside the scanner.** Participants repeated the picture-naming task outside the scanner, where their responses were recorded [1]. (*A*) Naming response time and (*B*) naming accuracy for Chinese and English on nonswitch and switch trials. (*C*) Overall naming accuracy across all trials. Dots are participants; violins show the distribution across participants.

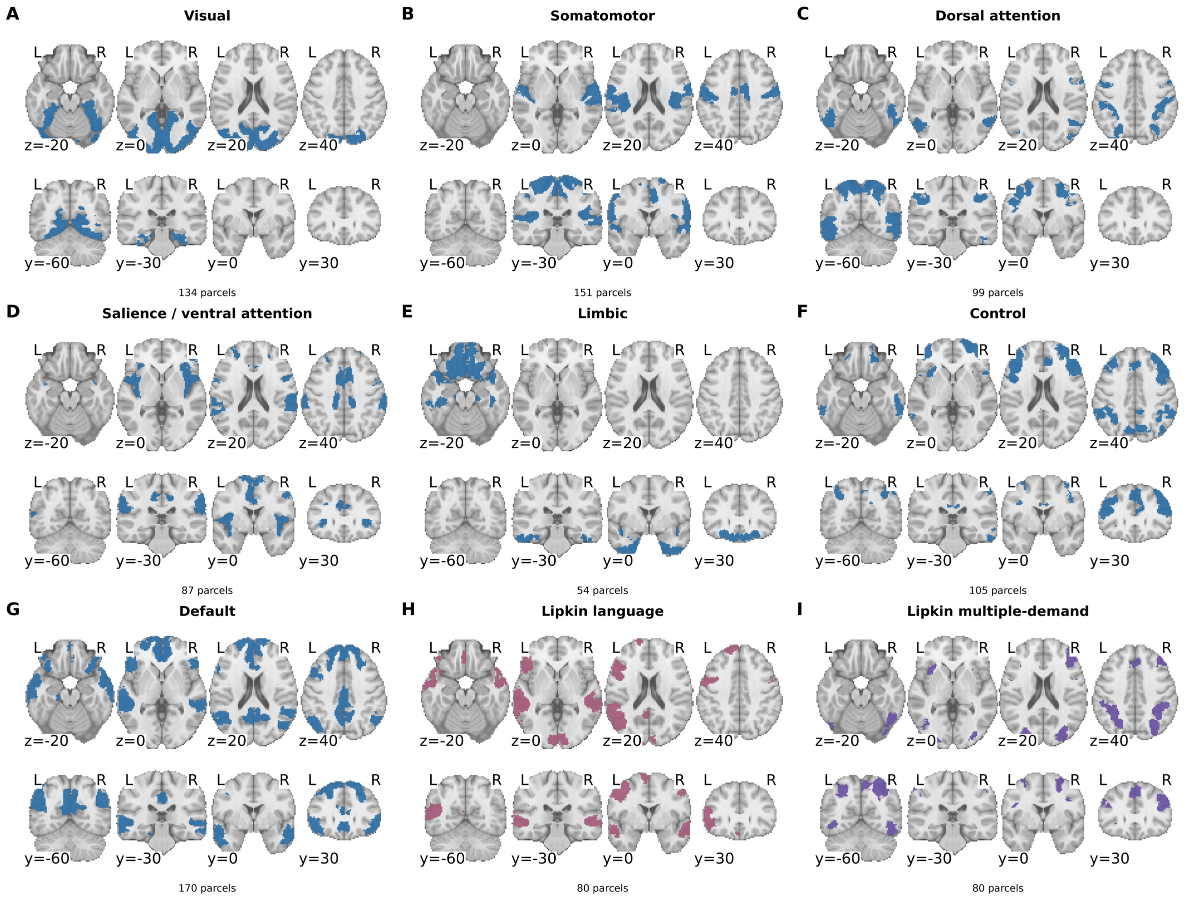


**Figure S12. Network and functional parcel sets.** (*A–G*) The seven Yeo networks [2] in Schaefer-800 space, with the number of parcels in each. (*H*) Language network and (*I*) multiple-demand network sets, each the 80 parcels (top 10%) with the highest mean probability in the corresponding probabilistic atlas [3, 4]. Axial ($z$) and coronal ($y$) slices are in MNI coordinates.

**Table S1. Within-person decoding and transfer between people, using the shared dimensions or the residual.** Balanced accuracy (%) across the 77 participants, given as mean ± SD and range (maximum minus minimum, percentage points). Within-person decoding is leave-one-run-out decoding within each participant. Learner transfer is, for each participant, the mean accuracy of the 76 classifiers trained on the other participants and tested on that participant. Teacher transfer is, for each participant, the mean accuracy of that participant's classifier tested on the 76 others. Learner and teacher transfer average over the same pairs, so their means are identical and only their spread differs. *Shared dimensions*: scores on the first $k$ shared dimensions. *Residual*: activity remaining after removing them. Chance is 50%.

| | | Within-person | | Learner transfer | | Teacher transfer | |
|---|---|---|---|---|---|---|---|
| $k$ | Representation | Mean ± SD | Range | Mean ± SD | Range | Mean ± SD | Range |
| 1 | Shared dimensions | 50.3 ± 4.2 | 18.1 | 50.3 ± 0.9 | 4.4 | 50.3 ± 1.2 | 2.7 |
| | Residual | 59.8 ± 5.7 | 28.1 | 57.1 ± 2.7 | 12.1 | 57.1 ± 2.0 | 11.3 |
| 5 | Shared dimensions | 54.7 ± 5.0 | 23.9 | 54.1 ± 2.9 | 13.5 | 54.1 ± 2.7 | 11.2 |
| | Residual | 59.8 ± 5.8 | 29.3 | 55.8 ± 2.2 | 10.9 | 55.8 ± 1.7 | 7.6 |
| 10 | Shared dimensions | 56.0 ± 5.6 | 29.1 | 55.1 ± 3.0 | 12.8 | 55.1 ± 2.2 | 10.2 |
| | Residual | 59.8 ± 5.7 | 26.6 | 55.1 ± 1.9 | 9.4 | 55.1 ± 1.5 | 7.2 |
| 25 | Shared dimensions | 56.6 ± 5.9 | 30.9 | 56.0 ± 2.9 | 13.6 | 56.0 ± 2.2 | 11.3 |
| | Residual | 60.3 ± 5.6 | 29.7 | 53.8 ± 1.6 | 7.4 | 53.8 ± 1.2 | 5.8 |
| 50 | Shared dimensions | 58.3 ± 5.9 | 27.2 | 57.2 ± 2.9 | 13.9 | 57.2 ± 2.0 | 11.4 |
| | Residual | 59.7 ± 5.4 | 28.2 | 51.9 ± 0.9 | 3.8 | 51.9 ± 0.7 | 3.7 |
| 100 | Shared dimensions | 58.5 ± 5.5 | 26.2 | 57.4 ± 2.7 | 13.1 | 57.4 ± 2.0 | 11.5 |
| | Residual | 60.1 ± 5.8 | 31.3 | 51.2 ± 0.7 | 2.7 | 51.2 ± 0.6 | 2.5 |
| 200 | Shared dimensions | 58.8 ± 5.4 | 27.8 | 57.2 ± 2.7 | 12.8 | 57.2 ± 1.9 | 10.2 |
| | Residual | 59.7 ± 5.9 | 28.4 | 50.9 ± 0.6 | 2.8 | 50.9 ± 0.6 | 2.9 |

**Table S2. Age of English acquisition is predicted after correction across all candidate measures.** Age of English acquisition (AoA) was predicted with leave-one-participant-out ridge regression from each of 43 candidate measures: learner transfer, teacher transfer and within-person decoding, each computed on the first $k$ shared dimensions or on the residual after their removal ($k = 1, 5, 10, 25, 50, 100, 200$), and within-person decoding on the full cortical pattern. To account for selecting the strongest measure, AoA values were permuted 10,000 times, the full search was repeated for each permutation, and the maximum cross-validated $R^2$ was retained to form the null distribution.

| Quantity | Value |
|---|---|
| Participants | 77 |
| Candidate measures | 43 |
| Permutations | 10,000 |
| Best measure | Learner transfer, first 10 shared dimensions |
| Pearson $r$ (observed vs. predicted AoA) | 0.441 |
| Cross-validated $R^2$ | 0.194 |
| 95th percentile of max-null $R^2$ | 0.070 |
| 99th percentile of max-null $R^2$ | 0.105 |
| Max-statistic corrected $p$ | 0.0003 |

**Table S3. Chronological age is predicted less well and does not survive correction.** Chronological age was predicted from the same 43 candidate measures with the same procedure as age of English acquisition (Table S2), with age values permuted 10,000 times.

| Quantity | Value |
|---|---|
| Participants | 77 |
| Candidate measures | 43 |
| Permutations | 10,000 |
| Best measure | Learner transfer, first 5 shared dimensions |
| Pearson $r$ (observed vs. predicted age) | 0.245 |
| Cross-validated $R^2$ | 0.055 |
| 95th percentile of max-null $R^2$ | 0.069 |
| 99th percentile of max-null $R^2$ | 0.115 |
| Max-statistic corrected $p$ | 0.086 |
